# Age-dependent PD-1 induction restricts IL-2-driven effector T cell responses during La Crosse virus infection in mice

**DOI:** 10.64898/2026.05.18.725972

**Authors:** Reem Alatrash, Sanjana Iyer, Bobby Brooke Herrera

**Author notes:** **Corresponding Author**: BBH.

## Abstract

Age is a major determinant of disease severity following La Crosse virus (LACV) infection, yet the immunological mechanisms underlying heightened susceptibility in children remains poorly defined. Here, we show that acute LACV infection in weanling mice induces T cell dysfunction characterized by early PD-1 upregulation and impaired effector differentiation despite evidence of activation. This state is associated with reduced IL-2-dependent STAT5 signaling, indicating a failure to respond to available cytokine cues. Although regulatory T cells expand and exhibit elevated CD25 expression, their depletion increases IL-2 levels without restoring antiviral T cell responses or viral control. In contrast, PD-1 blockade partially restores T cell activation, and combined PD-1 blockade with CD25 targeting enables robust effector differentiation and improved viral control. These findings demonstrate that checkpoint signaling limits T cell responsiveness to IL-2, uncoupling activation from differentiation and driving age-dependent susceptibility to LACV infection.

## Introduction

Age is a major determinant of disease severity in viral infections, yet the mechanisms underlying heightened susceptibility in children remain poorly defined. La Crosse virus (LACV), a mosquito-borne orthobunyavirus and the leading cause of pediatric arboviral encephalitis in the United States, represents an example of age-restricted neuroinvasive viral disease [1]. Infection is frequently asymptomatic or manifests as a mild febrile illness, but a subset of cases progress to severe disease characterized by seizures, encephalopathy, or death [2]. These outcomes occur almost exclusively in children, indicating that host age, rather than viral factors alone, governs disease progression [3, 4].

Murine models recapitulate this age-dependent phenotype. Weanling mice develop severe neurological disease following peripheral LACV infection, whereas adult mice are largely resistant to viral challenge [5, 6]. This resistance does not reflect differences in viral neurovirulence, as intracranial inoculation remains uniformly lethal across ages [7–10]. Rather, adult hosts effectively restrict viral dissemination to the central nervous system (CNS), whereas antiviral immunity in young hosts fails to do so.

Innate immune pathways play a critical role in early viral control. Type I interferon (IFN) signaling limits LACV replication and systemic spread, and disruption of this pathway increases susceptibility [8, 11]. Additional host factors, including MxA, MAVS, and Toll-like receptor signaling, further contribute to protection against neuropathology [12–14]. More recently, host determinants such as Connexin43 (Cx43/Gja1) and EphrinA2 (Efna2) have been implicated in regulating age-dependent susceptibility [15], underscoring the complexity of host-virus interactions governing viral neuroinvasion. These mechanisms, however, do not fully account for the failure of young hosts to control infection despite engagement of antiviral pathways.

The contribution of adaptive immunity to age-dependent protection remains less clearly defined. Leukocyte infiltration is observed in both human disease and experimental models [5, 16, 17], yet the functional consequences of these responses differ by age. CD4^+^ and CD8^+^ T cells are dispensable for CNS pathology in susceptible weanling mice but are required for protection in adult animals [9], indicating that the quality rather than the presence of adaptive responses determines outcome. Consistent with this, adult mice generate robust, polyfunctional antiviral T cell responses, whereas weanling mice mount weak responses and succumb to infection [18, 19]. Importantly, enhancing T cell immunity in young animals restores cytotoxic T cell function and improves survival, demonstrating that protective responses can be elicited when appropriately engaged [18].

These observations argue against a model of simple immune immaturity and instead point toward active regulation or dysregulation of antiviral responses in early life. The developing immune system is increasingly understood as a tightly controlled environment biased toward limiting immunopathology rather than maximizing effector function [20–22]. Such constraints raise the possibility that inhibitory pathways exert disproportionate influence over antiviral immunity in young hosts.

Programmed cell death protein 1 (PD-1) is a central inhibitory checkpoint receptor that constrains T cell activation, proliferation, and effector function [23–25]. Although most extensively studied in chronic infection and cancer, where it enforces T cell exhaustion, PD-1 is also rapidly induced during acute viral infection and can restrict effector differentiation even in the absence of persistent antigen presence [26–28]. However, the role of PD-1 in acute neurotropic infection, and its contribution to age-dependent susceptibility, remains incompletely defined.

Here, we demonstrate that acute LACV infection in weanling mice induces early and sustained PD-1 signaling that constrains antiviral T cell responses despite clear evidence of activation. This checkpoint engagement is associated with impaired interleukin-2 (IL-2)/signal transducer and activator of transcription 5 (STAT5) signaling and failure of effector differentiation, defining a state in which T cells are activated but unable to respond to essential cytokine cues. Restoration of antiviral immunity requires both relief of PD-1-mediated inhibition and modulation of the regulatory environment. These findings identify checkpoint signaling as a central determinant of age-dependent susceptibility to LACV infection and reveal a mechanism by which inhibitory pathways uncouple T cell activation from functional differentiation in early life.

## Results

### LACV infection induces age-dependent upregulation of T cell inhibitory receptors

To determine whether LACV infection is associated with T cell dysfunction, we examined expression of inhibitory receptors on CD4⁺ and CD8⁺ T cells at day 6 post-infection (dpi), a time point corresponding to advanced disease in weanling mice but preceding uniform mortality. We focused on PD-1, a well-established regulator of T cell activation and effector function and assessed additional inhibitory receptors including cytotoxic T-lymphocyte-associated protein 4 (CTLA-4), T cell immunoglobulin and mucin-domain containing-3 (TIM-3), and lymphocyte activation gene-3 (LAG-3). Comparisons were performed between weanling and adult mice under both uninfected and infected conditions.

At 6 dpi, LACV-infected weanling mice exhibited a significant increase in PD-1 expression on both CD4⁺ and CD8⁺ T cells compared to age-matched uninfected controls (Fig. 1A, 1E, S1). In contrast, PD-1 expression in adult mice remained unchanged following infection and comparable to uninfected controls (Fig. 1A, 1E, S1). Notably, PD-1 expression was significantly higher in infected weanling mice than in infected adults across both T cell compartments, indicating a pronounced age-dependent induction of inhibitory signaling.

**Figure 1.**
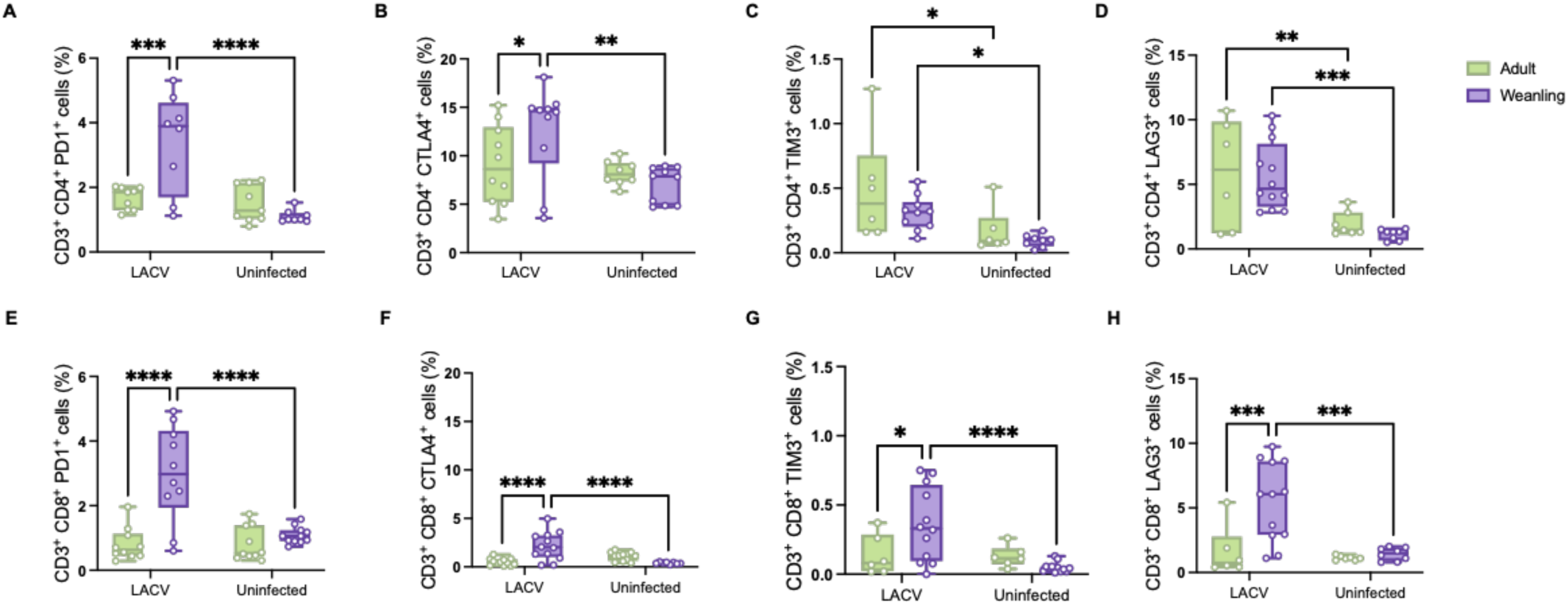
LACV infection induces age-dependent upregulation of inhibitory T cell pathways. (A, E) Frequencies of PD-1-expressing CD4⁺ and CD8⁺ T cells, (B, F) CTLA4-expressing CD4⁺ and CD8⁺ T cells, (C,G) TIM3-expressing CD4⁺ and CD8⁺ T cells and (D, H) LAG3-expressing CD4⁺ and CD8⁺ T cells in weanling and adult mice following LACV infection as assessed by flow cytometry at 6 dpi. Statistical significance was determined using one-way ANOVA test, ∗p < 0.05, ∗∗p < 0.01, ∗∗∗p < 0.001, ∗∗∗∗p < 0.0001. (n=6-13 in each group, data are representative of three independent experiments).

We next assessed whether additional inhibitory receptors followed similar patterns. CTLA-4 expression was significantly upregulated on CD4⁺ T cells from infected weanling mice compared to infected adults, with no differences in uninfected controls, indicating infection-dependent regulation (Fig. 1B, S2A). A similar trend was observed in CD8⁺ T cells (Fig.1F, S2B). TIM-3 expression increased following infection across age groups in CD4⁺ T cells but did not differ between ages (Fig.1C, S3A), whereas CD8⁺ T cells from infected weanling mice exhibited significantly higher TIM-3 expression compared to adults (Fig. 1G, S3B). LAG-3 expression increased on CD4⁺ T cells following infection independent of age (Fig. 1D, S4A), while CD8⁺ T cells expression displayed age-dependent differences (Fig. 1H, S4B).

Among these receptors, PD-1 displayed the most robust, consistent, and age-restricted upregulation, suggesting that it represents the dominant inhibitory axis distinguishing weanling from adult immune responses. We therefore focused subsequent analyses on the PD-1/PD-L1 pathway as a central candidate mechanism underlying impaired antiviral immunity in young hosts. Moreover, to determine whether this inhibitory signal was reinforced at the tissue level, we quantified expression of programmed death-ligand 1 (PD-L1). PD-L1 transcripts were significantly increased in the spleens of LACV-infected weanling mice compared to uninfected controls (Fig. S5B), with even higher expression observed in the brain, a primary site of viral replication and neuropathology (Fig. S5A). Additionally, infection of mouse fibroblasts in vitro resulted in a dose-dependent induction of PD-L1 expression (Fig. S5C), demonstrating that viral infection alone is sufficient to drive PD-L1 upregulation. Together, these data define an inhibitory landscape in weanling mice characterized by coordinated upregulation of PD-1 and other inhibitory receptors on T cells and PD-L1 within infected tissues, consistent with the establishment of a suppressive environment that may constrain antiviral T cell responses.

### LACV infection impairs T cell activation and differentiation in weanling mice

Given the elevated expression of inhibitory receptors, we next examined classical markers of T cell activation and differentiation. Expression of CD44 and CD62L on CD4⁺ and CD8⁺ T cells was assessed at 6 dpi to define naïve (CD44⁻CD62L⁺), effector (CD44⁺CD62L⁻), central memory (CD44⁺CD62L⁺), and intermediate populations. Adult mice mounted robust T cell responses following LACV infection, characterized by a significant increase in the frequency of effector CD4⁺ T cells compared to both uninfected controls and infected weanling mice (Fig. 2A, 2E, S6A). Effector CD4⁺ T cells were present in infected weanling mice; however, these cells exhibited significantly higher PD-1 expression compared to adult counterparts, indicating persistent inhibitory signaling despite acquisition of an effector phenotype (Fig. 2C, S6A). In the CD8⁺ T cell compartment, infected weanling mice displayed an increased frequency of effector CD8⁺ T cells compared to infected adults. Despite this apparent expansion, these cells expressed significantly higher levels of PD-1, suggesting that differentiation occurs in the presence of sustained inhibitory signaling (Fig. 2B, 2D, 2F, S6B).

**Figure 2.**
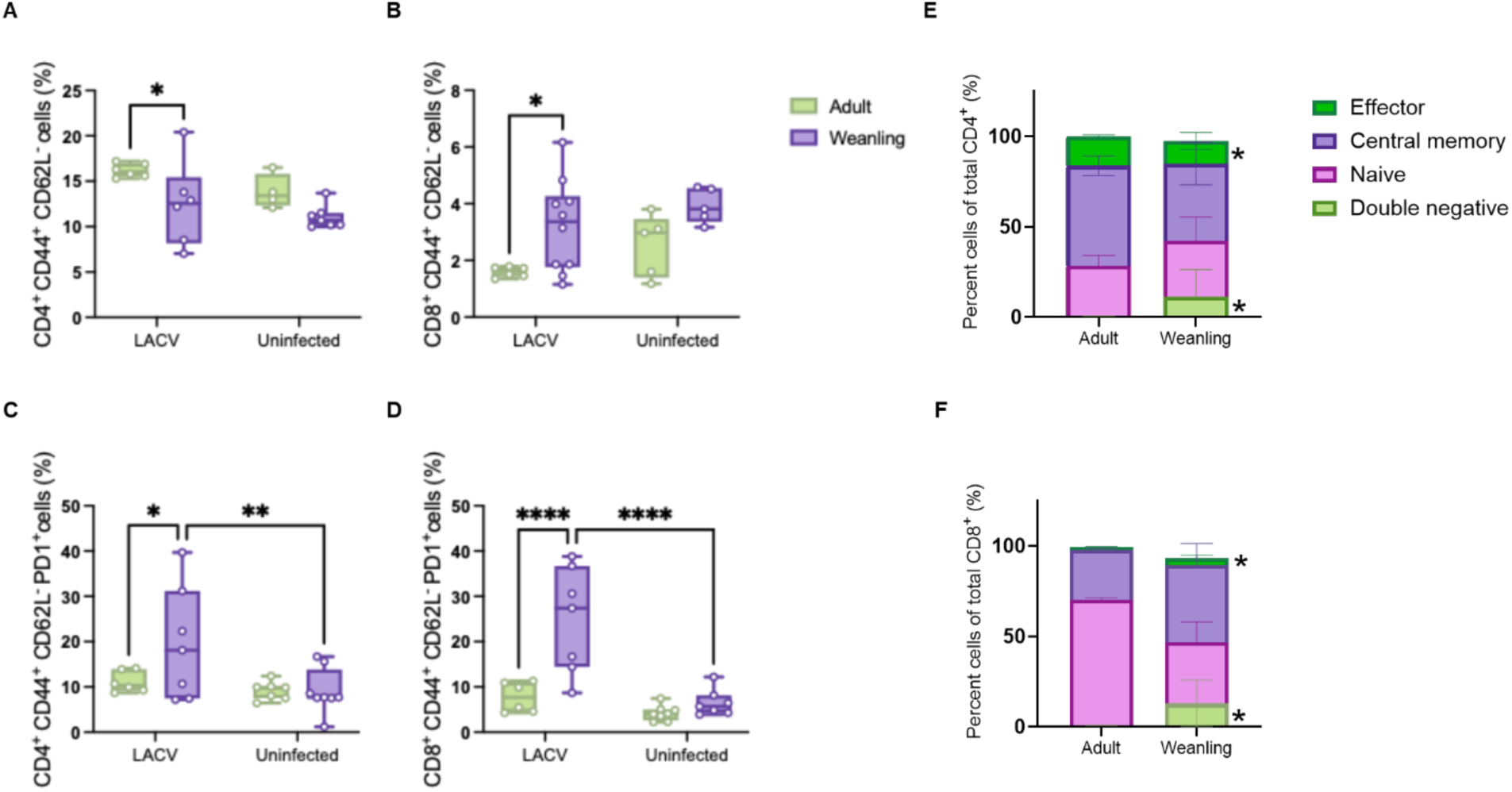
Age-dependent heterogeneity in T cell differentiation and activation during LACV infection. (A, B) Frequencies of effector CD4⁺ and CD8⁺ T cells, (C, D) Frequencies of PD-1-expressing effector CD4⁺ and CD8⁺ T cells. (E, F) Flow cytometry quantification of CD4⁺ and CD8⁺ T cells T cell subsets. (G) Representative flow cytometry plots LACV-infected adult and weanling mice demonstrating distinct T cell activation and differentiation states showing altered distribution and aberrant differentiation/activation profiles (as indicated by arrows) in weanling mice. Statistical significance was determined using one-way ANOVA test, ∗p < 0.05, ∗∗p < 0.01, ∗∗∗p < 0.001, ∗∗∗∗p < 0.0001. (n=6-10 in each group, data are a representative of three independent experiments).

In addition to these population-level differences, we observed a striking alteration in differentiation trajectories in a subset of infected weanling mice. Rather than accumulating conventional effector or memory populations, these animals exhibited a prominent CD44⁻CD62L^lo^ T cell population that was largely absent in adult mice (Fig. S7). This population was most pronounced within the CD4⁺ compartment and significantly enriched in infected weanling mice compared to age-matched uninfected controls (Fig. S7). The coexistence of CD62L downregulation, indicative of early activation, with failure to upregulate CD44 suggests that T cells initiate activation but do not complete the differentiation program. This pattern is consistent with a state of abortive or incomplete differentiation, in which T cells receive sufficient signals to exit quiescence but fail to acquire full effector functionality. Accordingly, infected weanling mice retain a substantial proportion of T cells in naïve or poorly differentiated states despite ongoing infection, in contrast to adult mice, in which T cells progress efficiently toward terminal effector phenotypes (Fig. 2). These findings demonstrate that LACV infection in weanling mice is associated with a qualitative defect in T cell differentiation, characterized not by absence of activation but by failure to transition into fully functional effector states in the presence of sustained inhibitory signaling.

### Altered IL-2/STAT5 signaling and expansion of a PD-1^hi^ regulatory T cell compartment in weanling mice

The observed defects in T cell differentiation prompted us to examine whether cytokine pathways required for effective T cell expansion and programming are differentially engaged. IL-2 plays a central role in promoting T cell proliferation, survival, and differentiation, while also maintaining regulatory T cell homeostasis. To assess IL-2-dependent signaling, we measured phosphorylation of STAT5 (pSTAT5) within CD4⁺ T cells at 6 dpi.

Adult mice exhibited significantly higher levels of STAT5 phosphorylation following infection, consistent with effective IL-2 receptor engagement and downstream signaling (Fig. 3A). In contrast, STAT5 activation was markedly reduced in weanling mice, indicating impaired IL-2 responsiveness despite ongoing infection. This difference suggests that defects in cytokine signaling may contribute directly to the failure of effector differentiation observed in young hosts.

**Figure 3.**
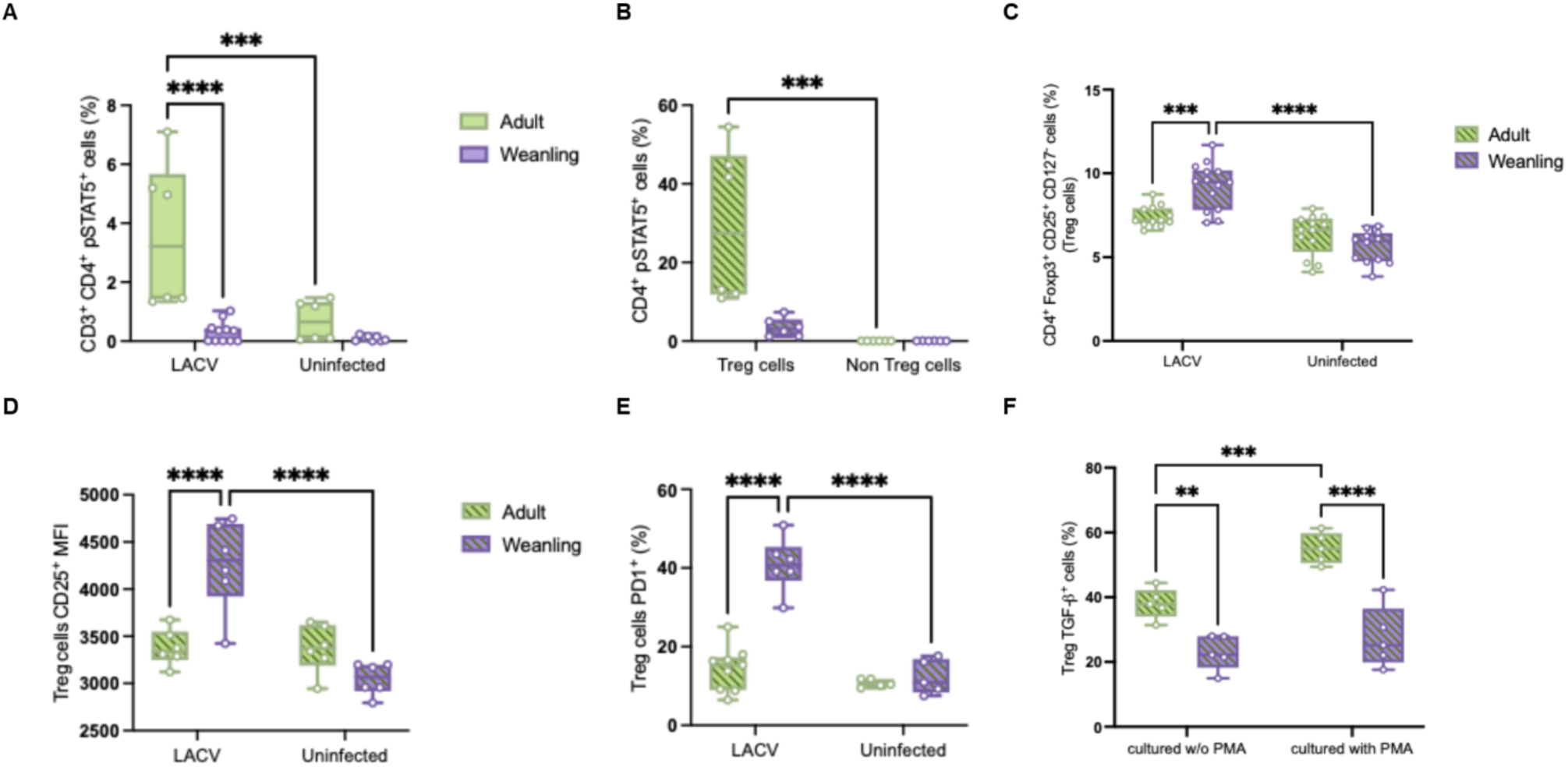
Altered IL-2/STAT5 signaling and expansion of a PD-1^hi^ regulatory T cell compartment in weanling mice following LACV infection. (A) Frequency of pSTAT5⁺ cells within the total CD4⁺ T cell compartment in weanling and adult mice following infection. (B) Distribution of pSTAT5 expression within Foxp3⁺ regulatory T cells (Tregs) and Foxp3⁻ conventional CD4⁺ T cells, demonstrating preferential STAT5 phosphorylation within the regulatory T cell compartment in both age groups. (C) Frequency of CD25⁺Foxp3⁺ CD4⁺ regulatory T cells in weanling and adult mice under naïve and LACV-infected conditions at day 6 post-infection. (D) Mean fluorescence intensity (MFI) of CD25 expression on Foxp3⁺ regulatory T cells. (E) Frequency of PD-1-expressing cells within the CD25⁺Foxp3⁺ regulatory T cell compartment in weanling and adult mice following infection. (F) TGF-β expression by regulatory T cells following ex vivo stimulation with Phorbol 12-myristate 13-acetate (PMA) compared to unstimulated controls in both age groups. Statistical significance was determined using one-way ANOVA test, ∗p < 0.05, ∗∗p < 0.01, ∗∗∗p < 0.001, ∗∗∗∗p < 0.0001. (n=3-15 in each group, data are a representative of three independent experiments).

Analysis of STAT5 distribution revealed that phosphorylation was enriched within the Foxp3^+^ compartment in both age groups (Fig. 3B, S8), reflecting preferential engagement of IL-2 signaling within regulatory T cells (Tregs). We therefore examined the abundance and phenotype of Tregs in greater detail. While Tregs were readily detected in peripheral tissues, they were rare or undetectable in the brain under both uninfected and infected conditions (Fig. S11), suggesting that their primary influence occurs within secondary lymphoid compartments rather than at sites of neuropathology.

LACV-infected weanling mice exhibited a significantly higher frequency of CD25⁺Foxp3⁺ CD4⁺ Tregs compared to infected adult mice and uninfected controls (Fig. 3C, S9). In addition to their increased abundance, Tregs from infected weanling mice displayed significantly higher CD25 expression, indicating increased expression of the high-affinity IL-2 receptor and suggesting enhanced capacity for IL-2 capture (Fig. 3D). Despite this, overall STAT5 phosphorylation within the CD4^+^ compartment remained low, indicating that increased receptor expression does not translate into effective downstream signaling.

Further phenotypic analysis revealed that Tregs from infected weanling mice expressed significantly higher levels of PD-1 compared to adult Tregs, consistent with altered regulatory programming (Fig. 3E, S10). Functional assessment demonstrated reduced TGF-β production following stimulation (Fig. 3F), indicating diminished functional responsiveness despite increased abundance. These findings define a state in which IL-2 signaling is both quantitatively and qualitatively altered in weanling mice. Expansion of a CD25^hi^ PD-1^hi^ Treg compartment occurs alongside impaired responsiveness rather than simple cytokine deprivation.

### Treg depletion alone is insufficient to restore antiviral T cell responses in weanling mice

The expansion of a PD-1^hi^ Treg compartment, together with impaired IL-2/STAT5 signaling, raised the possibility that Tregs might limit antiviral immunity through preferential consumption of IL-2 or direct suppression of effector responses. To test this, we depleted CD25⁺ cells in vivo and assessed the impact on viral control and T cell function. Efficient depletion of CD25⁺Foxp3⁺ Tregs was confirmed by flow cytometric analysis (Fig. S11), establishing effective disruption of this compartment prior to analysis.

Despite efficient depletion, viral burden in the brain was not reduced. Instead, anti-CD25-treated weanling mice exhibited modestly increased viral loads relative to untreated controls (Fig. 4A), indicating that removal of the regulatory compartment does not enhance control of LACV replication and may instead exacerbate viral dissemination. Consistent with this, depletion of CD25^+^ cells failed to improve T cell activation or differentiation (Fig. 4B-C). The frequency and distribution of CD44^+^CD62L^-^ effector CD4^+^ and CD8^+^ T cells remained unchanged, and no significant increase in activated T cell populations was observed (Fig. 4B-C). These findings indicate argue against a dominant role for Tregs as the primary constraint on antiviral T cell responses in this setting.

**Figure 4.**
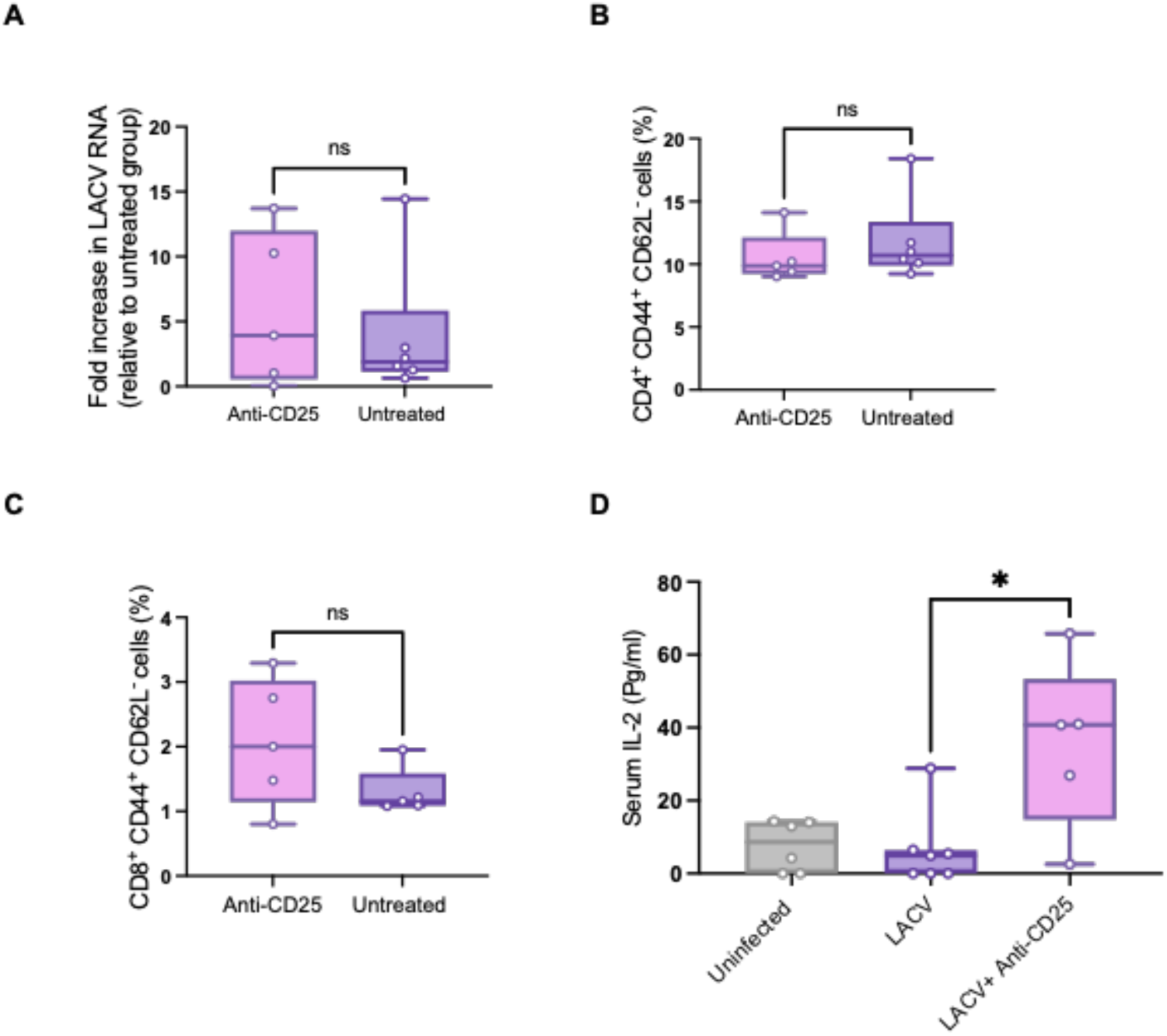
Regulatory T cell depletion alone is insufficient to restore antiviral T cell responses in weanling mice following LACV infection. (A) Viral burden in brain from uninfected and anti-CD25–treated weanling mice at day 6 post-infection, as measured by quantitative RT-PCR. (B) Frequency of activated/effector CD4⁺ and (C) CD8⁺ T cells following anti-CD25 treatment, assessed by CD44/CD62L expression. (D) Serum IL-2 concentrations in uninfected, LACV infected and anti-CD25-treated weanling mice at day 6 post-infection. Statistical significance was determined using Mann-Whitney U test, ∗p < 0.05, ∗∗p < 0.01, ∗∗∗p < 0.001, ∗∗∗∗p < 0.0001. (n=5-7 in each group, data are a representative of three independent experiments).

As expected, depletion of CD25^+^ cells results in increased circulating IL-2 levels (Fig. 4D), consistent with reduced cytokine consumption. However, this increase in IL-2 availability did not translate into enhanced STAT5 signaling, improved T cell differentiation, or reduced viral burden. The dissociation between cytokine availability and functional response suggest that impaired antiviral immunity in weanling mice cannot be explained solely by IL-2 deprivation or Treg-mediated suppression. Rather, these data point to a defect in the ability of T cells to respond to IL-2, even when it is present in excess. These findings demonstrate that depletion of Tregs and augmentation of IL-2 availability are insufficient to restore antiviral immunity in weanling mice, indicating that additional inhibitory mechanisms actively constrain T cell responsiveness during LACV infection.

### PD-1 blockade is required for restoration of effector T cell differentiation and viral control

The persistence of impaired T cell function despite increased IL-2 availability suggested that inhibitory signaling pathways may limit the ability of T cells to utilize cytokine cues. Given the pronounced upregulation of PD-1 observed in weanling mice, we next tested whether relief of PD-1-mediated inhibition is required to restore T cell responsiveness. As such, we evaluated the effects of PD-1 blockade alone or in combination with CD25 targeting on viral burden and T cell differentiation.

Blockade of PD-1 alone did not significantly reduce viral load compared to untreated controls (Fig. 5A), indicating that inhibition of this pathway in isolation is insufficient to restore antiviral control. In contrast, combined PD-1 blockade and CD25 targeting resulted in a significantly greater reduction in viral burden (Fig. 5A), demonstrating that relief of inhibitory signaling enables effective control of infection when coupled with modulation of the cytokine environment. This effect was accompanied by changes in systemic cytokine dynamics, as serum IL-2 levels increased following combined treatment but remained lower than those observed with CD25 targeting along (Fig. 5B), consistent with increased utilization of IL-2 under conditions of restored responsiveness.

**Figure 5.**
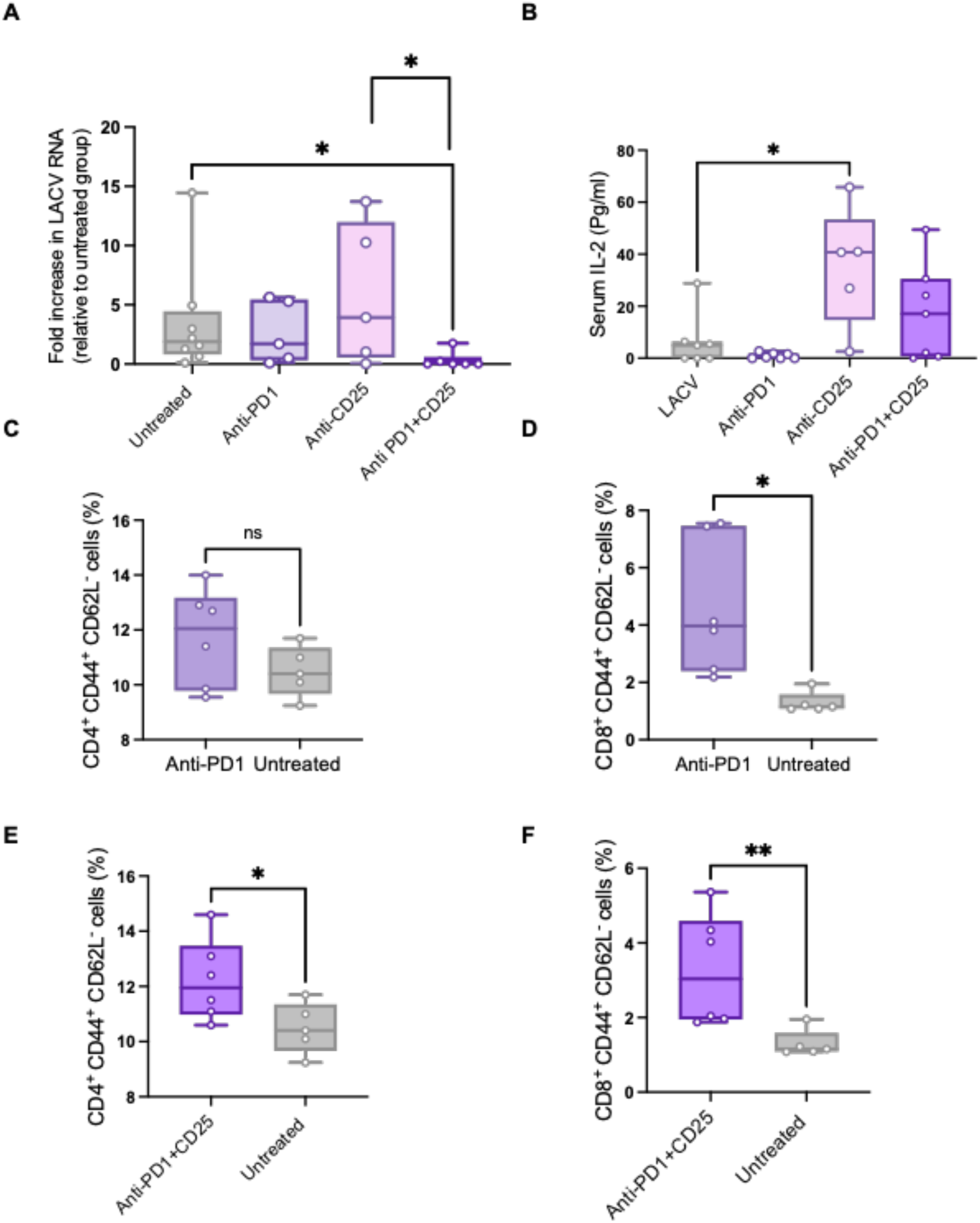
PD-1 blockade is required for restoration of effector T cell differentiation and viral control in weanling mice during LACV infection. (A) Brain viral burden in untreated weanling mice, anti-PD-1 antibody, anti-CD25 antibody, or combined anti-PD-1 and anti-CD25 antibodies, measured by quantitative RT-PCR at day 6 post-LACV infection. (B) Serum IL-2 concentrations in treated weanling mice at day 6 post-infection. (C–D) Frequency of effector CD4⁺ and CD8⁺ T cells, assessed by CD44/CD62L expression, following PD-1 blockade alone. (E–F) Frequency of effector CD4⁺ and CD8⁺T cells assessed by CD44/CD62L expression, following combined PD-1 and CD25 targeting. Statistical significance was determined using Mann-Whitney U test, ∗p < 0.05, ∗∗p < 0.01, ∗∗∗p < 0.001, ∗∗∗∗p < 0.0001. (n=5-6 in each group, data are a representative of three independent experiments).

Analysis of T cell responses revealed a similar pattern. PD-1 blockade alone produced a modest increase in T cell activation, reaching statistical significance within the CD8^+^ T cell compartment (Fig. 5D), but did not substantially alter overall differentiation. In contrast, combined PD-1 blockade and CD25 targeting resulted in a pronounced increase in activated T cells, including enhanced frequencies of effector CD4⁺ and CD8⁺ T cells (Fig. 5E-F). Notably, these effects were not observed following CD25 targeting in the absence of PD-1 blockade, reinforcing the conclusion that increased IL-2 availability alone is insufficient to drive effector differentiation (Fig. 5E-F). These findings establish a hierarchical relationship between cytokine signaling and checkpoint inhibition in which PD-1 constrains the capacity of T cells to respond to IL-2. Even in the presence of increased cytokine availability, effector differentiation cannot proceed unless inhibitory signaling is simultaneously relieved. Restoration of antiviral immunity therefore requires coordinated modulation of both inhibitory and regulatory pathways.

### Combined PD-1 blockade and CD25 targeting restore antigen-specific cytotoxic effector function of T cells

We next sought to determine whether restoration of T cell differentiation under combined PD-1 blockade and CD25 targeting translates into recovery of antigen-specific effector ceullar function. Splenocytes from treated and untreated weanling mice were stimulated ex vivo with LFn-LACV immunogens, and production of IFN-γ and granzyme B was assessed by intracellular cytokine staining.

Untreated weanling mice exhibited minimal antigen-specific IFN-γ production within the CD4^+^ T cell compartment following LFn-LACV stimulation (Fig. 6A). Combined PD-1 blockade and CD25 targeting resulted in a substantial increase in IFN-γ-producing CD4⁺ T cells, with the most pronounced responses observed against the LACV nucleoprotein (N) (Fig. 6A), indicating restoration of antigen-specific T cell function. In contrast, IFN-γ production by CD8⁺ T cells remained low in untreated mice and showed only a modest increase following combined treatment, which did not reach statistical significance (Fig. 6B). In comparison to IFN-γ, antigen-specific granzyme B responses were more robustly restored by treatment. CD4⁺ T cells stimulated with the glycoprotein Gc and N exhibited significantly increased granzyme B production in treated mice relative to untreated controls (Fig. 6C). Similarly, in the CD8⁺ T cell compartment, granzyme B production was reduced in untreated weanling mice but was strongly enhanced following combined PD-1 blockade and CD25 targeting (Fig. 6D), indicating recovery of cytotoxic effector capacity. These findings demonstrate that checkpoint-mediated inhibition not only limits T cell differentiation but also directly restricts acquisition of antigen-specific cytotoxic activity. Relief of PD-1 signaling, in conjunction with modulation of the IL-2 axis, enables functional reprogramming of antiviral T cells and restores effector capacity in young hosts.

**Figure 6.**
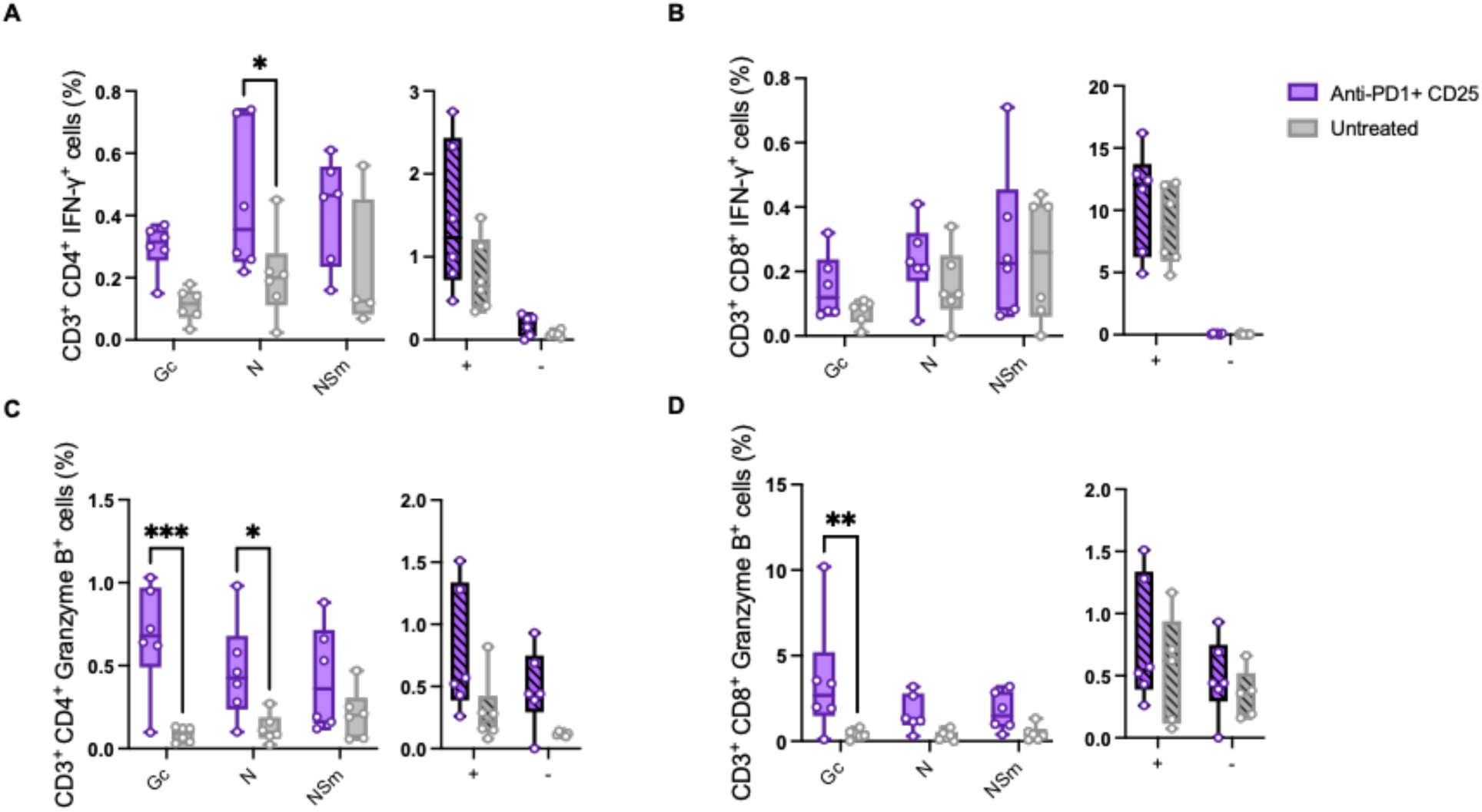
Combined PD-1 blockade and CD25 targeting restore antigen-specific cytotoxic effector function in weanling mice. Functional LACV-specific CD4^+^ and CD8^+^ T cell responses shown as the percent of CD4^+^ or CD8^+^ cells positive for IFN-γ (A and B) in CD3^+^ gate in flow cytometry/intracellular cytokine staining in treatment groups (A and B) and granzyme B (C and D) at 6 dpi. (−) negative control: cells stimulated with LFn alone; (+) positive control: cells stimulated with PMA (phorbol 12-myristate 13-acetate). Statistical significance was determined using ANOVA test, ∗*p* < 0.05, ∗∗*p* < 0.01, ∗∗∗*p* < 0.001, ∗∗∗∗*p* < 0.0001. Data are presented as individual points with minimum and maximum values, (*n* = 4-6 mice, data are combined from three independent experiments).

## Discussion

Age is a major determinant of disease severity following LACV infection, yet the mechanisms underlying heightened susceptibility in children have remained poorly understood. The present study demonstrates that acute LACV infection in weanling mice elicits a fundamentally distinct immune response characterized by early and sustained engagement of inhibitory pathways, impaired T cell differentiation, defective IL-2/STAT5 signaling, and failure to generate effective antigen-specific cytotoxic responses. Importantly, these defects occur despite clear evidence of T cell activation, indicating that antiviral immunity in young hosts is not absent, but actively constrained during a critical window of disease progression. These findings redefine age-dependent susceptibility to LACV infection as a consequence of dysregulated immune programming rather than simple developmental immaturity.

Among the inhibitory pathways, the PD-1/PD-L1 axis emerged as the dominant feature distinguishing susceptible weanling mice from resistant adults. Although PD-1 is most commonly associated with T cell exhaustion during chronic infection and cancer, increasing evidence indicates that this pathway is also rapidly engaged during acute viral infections, where it functions to temper effector responses and limit immunopathology [26, 29]. Early induction of PD-1 has been described in influenza virus, respiratory syncytial virus, West Nile virus, and acute lymphocytic choriomeningitis virus infection, where excessive checkpoint engagement can impair differentiation and functional maturation even in the absence of persistent antigen presence [30–35]. Similar observations have recently been reported in severe fever with thrombocytopenia syndrome virus (SFTSV), a distantly related bunyavirus, in which elevated PD-1/PD-L1 signaling correlates with impaired antiviral immunity [36]. The present findings extend this framework by demonstrating that checkpoint signaling can function as a major determinant of age-dependent susceptibility during acute neurotropic viral infection.

PD-1 was robustly induced on both CD4⁺ and CD8⁺ T cells specifically in weanling mice, indicating preferential engagement of inhibitory signaling in young hosts. This distinction was not simply quantitative but qualitatively associated with failed effector differentiation and impaired IL-2 responsiveness. While transient PD-1 induction is a normal component of early T cell activation, sustained expression in weanling mice was accompanied by defective STAT5 phosphorylation and accumulation of poorly differentiated T cell populations, suggesting that checkpoint signaling diverts antiviral responses away from productive effector programming. Mechanistically, PD-1 signaling attenuates proximal T cell receptor signaling, suppresses IL-2 production, and limits downstream pathways required for proliferation and differentiation [37–39]. The convergence of elevated PD-1 expression with impaired IL-2/STAT5 signaling observed here provides a mechanistic link between checkpoint engagement and defective antiviral immunity in early life.

Inhibitory signaling was further reinforced at the tissue level through increased PD-L1 expression in both lymphoid organs (e.g., spleen) and the brain, the principal site of viral replication and neuropathology. Similar spatial organization of PD-1/PD-L1 signaling has been described in neurotropic infections, where PD-L1 expression within the CNS limits T cell effector function and viral clearance [40, 41]. Several viruses actively induce PD-L1 expression in infected cells as a mechanism of immune modulation. Hantaviruses induce PD-L1 and PD-L2 expression in endothelial and dendritic cells [42], while Zika virus transcriptionally activates PD-L1 through viral protein-mediated mechanisms [43]. Consistent with these observations, LACV infection directly induced PD-L1 expression in fibroblasts in vitro indicating that viral infection alone is sufficient to establish a suppressive signaling environment independent of broader inflammatory cues. These findings suggest that PD-1 signaling in weanling mice is reinforced simultaneously at both the cellular and tissue levels, creating conditions that broadly constrain antiviral T cell function.

A striking feature of the weanling response was the uncoupling of T cell activation from terminal differentiation. Despite evidence of activation and expansion, T cells from infected weanling mice failed to efficiently transition into fully differentiated effector states and instead accumulated atypical CD44⁻CD62L^lo^ populations suggestive of incomplete or abortive differentiation. Similar dysfunctional trajectories have been described in settings of excessive checkpoint signaling, altered cytokine availability, or persistent antigen exposure, where T cells initiate activation programs but fail to acquire stable effector functionality [44–47]. Comparable intermediate or “transitory” dysfunctional states have been described during chronic viral infection and tumor-associated exhaustion programs [48–50]. The phenotype observed during acute LACV infection therefore suggests that inhibitory programming can emerge early during infection in susceptible hosts and may alter developmental trajectories of antiviral T cells before stable effector populations are established.

The altered cytokine landscape observed in weanling mice further supports this interpretation. Early-life immunity is increasingly recognized as a tightly regulated developmental state biased toward limiting immunopathology rather than maximizing inflammatory responses [51–53]. Tregs are central to this balance and exhibit heightened sensitivity to IL-2-dependent STAT5 signaling during early immune development [53–55]. Consistent with this biology, LACV-infected weanling mice exhibited expansion of a CD25⁺Foxp3⁺ Treg compartment together with preferential STAT5 phosphorylation within Foxp3^+^ cells [56]. However, despite elevated CD25 expression, global STAT5 activation remained markedly impaired, indicating that increased receptor abundance does not translate into effective downstream signaling. Tregs from weanling mice also expressed elevated PD-1 and exhibited diminished TGF-β production following stimulation, suggesting altered functional programming rather than enhanced suppressive capacity. PD-1 signaling has previously been shown to destabilize or restrain Treg function under specific inflammatory conditions [57], and Tregs can exhibit substantial phenotypic plasticity during disrupted cytokine signaling states [58]. These findings argue against a model of excessive immune suppression alone and instead support broader dysregulation of both regulatory and effector immune compartments.

One of the central findings of this study is that cytokine availability and cytokine responsiveness are mechanistically distinct. Although Tregs are recognized as dominant consumers of IL-2 [56], depletion of CD25^+^ cells increased circulating IL-2 levels without restoring effector differentiation or viral control. These findings indicate that defective antiviral immunity in weanling mice cannot be explained solely by cytokine deprivation. Rather, PD-1 signaling appears to limit the ability of T cells to respond to available IL-2, effectively uncoupling cytokine sensing from downstream differentiation programs. This distinction is particularly important because it reframes immune dysfunction during acute infection not as a failure to generate activating signals, but as a failure of T cells to appropriately interpret and utilize them.

Direct blockade of PD-1 blockade revealed the dominant role of checkpoint signaling in shaping immune outcomes during LACV infection. PD-1 blockade alone partially improved T cell activation but was insufficient to fully restore viral control, whereas combined PD-1 blockade and CD25 targeting resulted in robust effector differentiation, enhanced cytotoxic function, and significantly improved control of viral replication. These findings establish a hierarchical relationship between checkpoint signaling and cytokine responsiveness in which inhibitory signaling remains dominant even in cytokine-replete conditions. The efficacy of PD-1 blockade has similarly been linked to the balance between effector and Treg populations in cancer and chronic infection models [59], suggesting that restoration of immunity depends not only on relieving signaling but also on reconfiguring the broader regulatory environment.

These findings also extend broader concepts derived from chronic viral infection into the setting of acute pediatric viral disease. PD-1 blockade restores exhausted CD8^+^ T cell function during chronic LCMV infection [39, 60], and increasing evidence indicates that checkpoint pathways can similarly constrain immunity during acute infection [61]. The present study demonstrates that such mechanisms may be particularly consequential during early life, where developmental biases toward immune regulation amplify the impact of inhibitory signaling. Under these conditions, PD-1 functions not merely as a marker of activation or exhaustion, but as a central bottleneck governing the ability of T cells to undergo productive differentiation and acquire antiviral effector function.

Several limitations should be considered. First, analysis focused on a single disease-relevant time point and therefore does not fully capture the temporal evolution of checkpoint signaling and T cell differentiation earlier during infection. Second, the upstream mechanisms driving preferential PD-1/PD-L1 induction in weanling mice remain unresolved, and future studies are expected to determine whether they involve developmental differences in antigen presentation, innate sensing, or inflammatory cytokine production. Finally, while combined PD-1 blockade and CD25 targeting restored antiviral immunity, the potential impact of these interventions on immunopathology within the CNS and outcomes was not directly assessed, although remains an area of active investigation.

Taken together, these findings establish checkpoint-mediated immune regulation as a central determinant of age-dependent susceptibility to LACV infection. More broadly, they demonstrate that inhibitory pathways classically associated with chronic infection and cancer can shape immune outcomes during acute viral infection in specific developmental contexts. The developing immune system may therefore be uniquely vulnerable to checkpoint-mediated dysregulation, with important implications for pediatric neurotropic viral diseases beyond LACV.

## Resource Availability

### Lead contact

Further information and requests for resources and reagents should be directed to and will be fulfilled by the lead contact, Bobby Brooke Herrera.

### Materials availability

This study did not generate new unique reagents.

### Data and code availability

- All data supporting the findings of this study are available within the article and its supplemental information. Raw flow cytometry files and RT-qPCR results are available from the lead contact upon reasonable request.
- This study did not generate any original code.
- No additional resources were generated or analyzed during this study.

## Acknowledgments

We would like to thank Rutgers Global Health Institute, Rutgers Robert Wood Johnson Medical School, and the Child Health Institute of New Jersey for their support. We would also like to thank the Foundation for Health Advancement for grant support (IFHA 17-24). The funders had no role in the design, data collection, data analysis, and reporting of this study.

## Author contributions

Conceptualization, R.A. and B.B.H.; formal analysis, R.A. and B.B.H.; investigation, R.A., S.I. and B.B.H.; resources, B.B.H.; data curation, R.A.; writing – original draft preparation, R.A. and B.B.H.; writing – review and editing, R.A. and B. B.H.; visualization, R.A. and B.B.H.; supervision, B.B.H.; funding acquisition, B.B.H. All authors have read and agreed to the published version of the manuscript.

## Supplemental information

See document S1: Figures S1-S11.

## STAR★METHODS

### KEY RESOURCES TABLE

| REAGENT OR RESOURCE | SOURCE | IDENTIFIER |
| --- | --- | --- |
| Antibodies |  |  |
| Rat anti-mouse Granzyme B Monoclonal Antibody (NGZB), PE-Cyanine5.5 | Thermo Fisher Scientific | Cat# 35-8898-80, RRID:AB_2848328 |
| Rat anti-mouse IFN gamma Monoclonal Antibody (XMG1.2), eFluor™ 660 | Thermo Fisher Scientific | Cat# 50-7311-80, RRID:AB_11218909 |
| Rat anti-mouse CD3 Monoclonal Antibody (OKT3), Alexa Fluor™ 700, eBioscience | Thermo Fisher Scientific | Cat# 56-0037-42, RRID:AB_10714978 |
| Rat anti-mouse CD25 Monoclonal Antibody (PC61.5), eFluor™ 506, eBioscience™ | Thermo Fisher Scientific | Cat# 69-0251-82, RRID:AB_2762802 |
| Rat anti-mouse FOXP3 Monoclonal Antibody (FJK-16s), PE-Cyanine7, eBioscience™ | Thermo Fisher Scientific | Cat# 25-5773-82, RRID:AB_891552 |
| Rat anti-mouse CD127 Monoclonal Antibody (A7R34), Brilliant Violet™ 421, eBioscience™ | Thermo Fisher Scientific | Cat# 404-1271-82, RRID:AB_2929074 |
| Rat anti-mouse TGF Beta 1 Recombinant Rabbit Monoclonal Antibody (230544B7), PE | Thermo Fisher Scientific | Cat# 81746-2, RRID:AB_3670503 |
| Rat anti-mouse Phospho-STAT5 (Tyr694) Monoclonal Antibody (SRBCZX), PE, eBioscience™ | Thermo Fisher Scientific | Cat# 12-9010-42, RRID:AB_2572671 |
| Rat anti-mouse CD279 (PD-1) Monoclonal Antibody (J43), Super Bright™ 780, eBioscience™ | Thermo Fisher Scientific | Cat# 78-9985-82, RRID:AB_2734947 |
| Rat anti-mouse CD152 (CTLA-4) Monoclonal Antibody (UC10-4B9), APC, eBioscience™ | Thermo Fisher Scientific | Cat# 17-1522-82, RRID:AB_2016700 |
| Rat anti-mouse CD366 (TIM3) Monoclonal Antibody (RMT3-23), PE, eBioscience™ | Thermo Fisher Scientific | Cat# 12-5870-82, RRID:AB_465974 |
| Rat anti-mouse CD223 (LAG-3) Monoclonal Antibody (eBioC9B7W (C9B7W)), PE-Cyanine7, eBioscience™ | Thermo Fisher Scientific | Cat# 25-2231-82, RRID:AB_2573428 |
| Rat anti-mouse CD44 Monoclonal Antibody (IM7), Brilliant Violet™ 421, eBioscience™ | Thermo Fisher Scientific | Cat# 404-0441-82, RRID:AB_2925505 |
| Rat anti-mouse CD62L (L-Selectin) Monoclonal Antibody (MEL-14), Super Bright™ 600, eBioscience™ | Thermo Fisher Scientific | Cat# 63-0621-82, RRID:AB_2637416 |
| Rat anti-mouse CD4 Monoclonal Antibody (GK1.5), APC-eFluor™ 780 | Thermo Fisher Scientific | Cat# 47-0041-80, RRID:AB_11219883 |
| Rat anti-mouse CD8 Monoclonal Antibody (53-6.7) Alexa Fluor 488 | Thermo Fisher Scientific | Cat# 53-008180 |
| Anti-mouse CD16/CD32 Fcγ III/II (Thermo fisher sciences) | Thermo Fisher Scientific | Cat # 14-0161-82 |
| InVivoMab anti-mouse CD25 (IL-2Rα), Clone PC-61.5.3 | Bio X Cell | Cat # BE0012<br>RRID:AB_1107619 |
| InVivoMab anti-mouse PD-1 (CD279), Clone RMP1-14 | Bio X Cell | Cat # BE0146<br>RRID:AB_10949053 |
| Bacterial and virus strains |  |  |
| NIH: La Crosse Virus, NR-540 | Human isolate, BEI Resources | Catalog No. NR-540<br>(Derived from ATCC® VR-744™) |
| Biological samples |  |  |
| N/A |  |  |
| Chemicals, peptides, and recombinant proteins |  |  |
| LFn-LACV immunogens | Mir Biosciences, Inc | N/A |
| Brefeldin A Solution | Thermo Fisher Scientific | Cat# 00-4506-51 |
| RBC Lysis Buffer | Thermo Fisher Scientific | Cat# J62150.AP |
| eBioscience™ Foxp3 / Transcription Factor Staining Buffer Set | Thermo Fisher Scientific | Cat# 00-5523-00 |
| Intracellular Fixation & Permeabilization Buffer Set | Thermo Fisher Scientific | Cat# 88-8824-00 |
| Cell Stimulation Cocktail (plus protein transport inhibitors) | Thermo Fisher Scientific | Cat# 00-4975-93 |
| Critical commercial assays |  |  |
| Mouse IL-2 ELISA Kit | Thermo Fisher Scientific | Cat# BMS601 |
| LIVE/DEAD™ Fixable Red Dead Cell Stain Kit | Thermo Fisher Scientific | Cat# L34972 |
| RNeasy Lipid Tissue Mini Kit | Qiagen | Cat# 74804 |
| High-Capacity cDNA Reverse Transcription Kit | Thermo Fisher Scientific | Cat# 4368814 |
| PureLink™ RNA Mini Kit | Thermo Fisher Scientific | Cat# 12183018A |
| PowerTrack SYBR Green Master Mix | Applied Biosystems | Cat# A46109 |
| Deposited data |  |  |
| N/A |  |  |
| Experimental models: Cell lines |  |  |
| Vero cells | ATCC | Cat# CCL-81.4,<br>RRID:CVCL_0059 |
| Mouse Fibroblasts | ATCC | Cat# CRL-1658 |
| Experimental models: Organisms/strains |  |  |
| Mice C57BL/6 | The Jackson Laboratory | RRID:MGI:2159769 |
| Oligonucleotides |  |  |
| N/A |  |  |
| Recombinant DNA |  |  |
| N/A |  |  |
| Software and algorithms |  |  |
| Cytek Aurora Spectral Analyzer | Cytek | RRID:SCR_019826 |
| Cytek SpectroFlo | Cytek | RRID:SCR_025494 |
| FLOWJo version 10.2 | TreeStar, Inc | RRID:SCR_008520 |
| GraphPad Prism | GraphPad software | RRID:SCR_002798 |
| BioTek Synergy LX Multi-Mode Microplate Reader |  | RRID:SCR_019763 |
| QuantStudio Real-Time PCR Software | Applied Biosystems | SCR_021096 |
| BioRender | BioRender | RRID:SCR_018361 |
| Other |  |  |
| N/A |  |  |

### Methods

#### Experimental model

All animal experiments were approved by the Rutgers University Institutional Animal Care and Use Committee (IACUC, protocol number 202300007) and were conducted in accordance with institutional and national guidelines for the care and use of laboratory animals. Wild-type C57BL/6 mice were purchased from Jackson Laboratories and maintained in a specific pathogen-free breeding colony at the Child Health Institute of New Jersey. Both male and female mice were used in the study. Mice were age matched and used at either 3 weeks old (weanling stage) or 8 weeks old (adult stage) to model age-dependent susceptibility to La Crosse virus (LACV). No sex-specific differences were observed in any of the experiments, and results were pooled across sexes for all analyses. Where relevant, sex was balanced across experimental groups.

#### Infection of mice with LACV and neurological disease progression criteria

Wild-type (C57Bl/6) mice were purchased from Jackson Laboratories and maintained in a breeding colony at the Child Health Institute of New Jersey. LACV (La Crosse Virus, NR-540), a human isolate was purchased and obtained from BEI Resources. Mice at 3 (weanling) or 8 (adult) weeks of age were inoculated with 10^3^ PFU of LACV in phosphate-buffered saline (PBS) intraperitoneally in a volume of 200 μL/mouse. Mice were observed daily for signs of neurological disease that included hunched posture, seizures, reluctance, or inability to move normally, or paralysis. Animals with clear clinical signs of neurological disease were scored as clinical and euthanized immediately. As previously described, 100% of weanling mice were clinical at day 7 post infection.

#### Splenic leukocytes harvest

Spleens from LACV-infected and uninfected mice were harvested at 6 dpi, as described previously [18]. Spleens were processed into single-cell suspensions by passing them through a 70 μM mesh filter, followed by red blood cell lysis using RBC Lysis Buffer (Thermo Fisher Scientific). Isolated cells were then seeded for subsequent flow cytometry/ICS assays.

#### Flow cytometry and ICS assay

Briefly, isolated cells are seeded at a density of 2 x 10^6^ cells per well in a 96-well U-bottom plate. cells are stained for viability using a viability dye and Fc receptors are blocked using anti-mouse CD16/CD32 Fcγ III/II (ThermoFisher sciences) and normal mouse serum. Cells are then stained for extracellular markers using a panel of antibodies; (all from Thermo Fisher Scientific), washed, and fixed and permeabilized using the FIX & PERM Cell Fixation and Permeabilization Kit (ThermoFisher Scientific). Following permeabilization, cells are stained for intracellular cytokines; (ThermoFisher Scientific), washed, and resuspended in FACS buffer. Flow cytometry data are acquired using the Aurora system (Cytek Biosciences) and analyzed using FlowJo software (version 10.2; TreeStar, Inc), retaining live cells and excluding doublets using SSC-A and FSC-A gating.

#### LFn-LACV immunogens

LFn-LACV immunogens were purchased from Mir Biosciences, Inc.. In brief, gene fragments encoding the LACV Gc, Gn, N, NSm, NSs and RdRp were cloned into the LFn expression vector. The constructs were transformed into Escherichia coli BLR (DE3) (MilliporeSigma). Selected transformants were sequenced to verify in-frame cloning. The LFn-LACV immunogens were expressed upon induction with isopropyl-β-d-thiogalactopyranoside (IPTG; MilliporeSigma) in 5 liters of Luria broth containing Kanamycin for 4 h. Cells were pelleted by centrifugation and resuspended in imidazole (1 mM) binding buffer (Novagen,) in the presence of a protease inhibitor cocktail (ThermoFisher Scientific). Cell pellets were sonicated and centrifuged at 4°C, and the supernatants were loaded in an equilibrated nickel-charged column for affinity purification. The bound proteins were eluted in 100 to 200 mM imidazole, desalted with a Sephadex G-25M column (Sigma-Aldrich, St. Louis, MO, USA), and eluted in phosphate-buffered saline (PBS) (Sigma-Aldrich, St. Louis, MO, USA). The PBS-eluted proteins were passed through Detoxi-Gel (ThermoFisher Scientific). Protein concentrations were determined, and samples were stored at −80°C.

#### Quantification of viral RNA in brain tissue

Total RNA was extracted from homogenized mouse brains using the RNeasy Lipid Tissue Mini Kit (Qiagen), following the manufacturer’s protocol. Briefly, 20–30 mg of brain tissue was homogenized in QIAzol lysis reagent, followed by phase separation with chloroform and purification on silica-membrane spin columns. RNA concentration and purity were assessed using a NanoDrop One spectrophotometer (ThermoFisher Scientific). One microgram of total RNA was reverse transcribed into cDNA using the High-Capacity cDNA Reverse Transcription Kit (ThermoFisher Scientific) with random primers, in accordance with the manufacturer’s instructions.

Quantitative PCR (qPCR) was performed using the PowerTrack SYBR Green Master Mix (ThermoFisher Scientific) on a QuantStudio 3Flex Real-Time PCR System (Applied Biosystems). The following LACV-specific primers targeting the S segment were used: forward 5′-ATTCTACCCGCTGACCATTG-3′ and reverse 5′-GTGAGAGTGCCATAGCGTTG-3′ [15]. GAPDH was used as the internal control gene. The ΔCt was determined by subtracting the average GAPDH Ct from the corresponding LACV Ct. To evaluate relative viral load, ΔΔCt values were computed using the vaccinated group as the calibrator. Fold changes in viral RNA abundance relative to the vaccinated group were calculated using the 2^(-ΔΔCt) method.

#### Quantification of PD-L1 expression in spleen and brain tissue

Total RNA was extracted from homogenized spleen and brain tissues as described above for viral RNA quantification. RNA concentration and purity were measured using a NanoDrop One spectrophotometer (ThermoFisher Scientific), and 1 µg of total RNA was reverse transcribed into cDNA using the High-Capacity cDNA Reverse Transcription Kit (Thermo Fisher Scientific, Cat# 4368814) according to the manufacturer’s instructions. Quantitative PCR (qPCR) was performed using PowerTrack SYBR Green Master Mix (ThermoFisher Scientific) on a QuantStudio 3Flex Real-Time PCR System (Applied Biosystems). Mouse PD-L1 (Cd274) expression was measured using the following primers: forward 5′-TGCGGACTACAAGCGAATCACG-3′ and reverse 5′-CTCAGCTTCTGGATAACCCTCG-3′. GAPDH was used as the internal reference gene. Relative PD-L1 expression levels were calculated using the comparative Ct (ΔΔCt) method. ΔCt values were determined by subtracting the Ct value of GAPDH from the Ct value of PD-L1 for each sample. Fold changes in PD-L1 expression were calculated using the 2^(-ΔΔCt) method relative to the designated control group.

#### Quantitative PCR analysis of PD-L1 expression in LACV-infected fibroblasts

Mouse fibroblasts were seeded and infected with LACV at multiplicities of infection (MOI) of 0.1 or 1. At 72 hours’ time point post-infection, total RNA was extracted using RNA extraction kit according to the manufacturer’s instructions (ThermoFisher Scientific). RNA concentration and purity were assessed prior to downstream applications. Complementary DNA (cDNA) was synthesized from extracted RNA using a standard reverse transcription protocol.

Quantitative PCR (qPCR) was performed using a two-step SYBR Green-based detection method (as described above). Gene expression levels of *Cd274* (PD-L1) and LACV were quantified using gene-specific primers, with *Gapdh* serving as the endogenous control. Relative gene expression was calculated using the ΔΔCt method, normalizing target gene expression to *Gapdh* and comparing infected samples to mock-infected controls.

#### In vivo antibody blockade experiments

For in vivo depletion and checkpoint blockade experiments, monoclonal antibodies targeting CD25 and PD-1 were obtained from Bio X Cell (InVivoMAb anti-mouse CD25 [IL-2Rα], clone PC61, Cat# BE0012; InVivoMAb anti-mouse PD-1 [CD279], Cat# BE0146). To deplete Tregs, mice received 100 µg of anti-CD25 antibody via intraperitoneal (i.p.) injection on days −2 and −1 prior to LACV challenge. Efficient depletion of Tregs was confirmed by flow cytometric analysis starting 2 dpi. For PD-1 blockade experiments, mice were administered 100 µg of anti-PD-1 antibody via i.p. injection on days −2 and −1 prior to LACV infection, followed by a third dose at 3 dpi. Mice were euthanized at 6 dpi, and tissues were collected for downstream immunological analyses.

#### Serum IL-2 Quantification by ELISA

Serum interleukin-2 (IL-2) levels were quantified using a commercially available mouse IL-2 ELISA kit (ThermoFisher Scientific, Cat. No. BMS601) according to the manufacturer’s instructions. Briefly, serum samples were collected from mice at the indicated time points and stored at −80 °C until analysis. Samples and standards were added to 96-well plates pre-coated with anti-mouse IL-2 capture antibody and incubated as recommended. Following washing steps, a biotin-conjugated detection antibody was applied, followed by streptavidin–horseradish peroxidase (HRP). Signal was developed using tetramethylbenzidine (TMB) substrate and stopped with acidic stop solution. Absorbance was measured at 450 nm using a microplate reader, and IL-2 concentrations were calculated from a standard curve generated using recombinant IL-2 standards.

#### Quantification and statistical analysis

All statistical analyses were performed using GraphPad Prism v7.01. Flow cytometry data were analyzed using FlowJo v10.2. Statistical comparisons were made using one-way or two-way ANOVA with appropriate post hoc tests (as specified in figure legends) or The Mann-Whitney U test. For all analyses, p < 0.05 was considered statistically significant. Exact values of n (indicated in each figure legend) refer to the number of biological replicates, typically the number of individual mice per group. Data are presented as either mean ± standard deviation (SD) or individual data points with mean and range, as indicated. All statistical details including test used, definition of center, dispersion, p-values, and sample sizes can be found in the corresponding figure legends, figures, and results section of the manuscript.

