## Supplementary Information for "Age-dependent PD-1 induction restricts IL-2-driven effector T cell responses during La Crosse virus infection in mice"

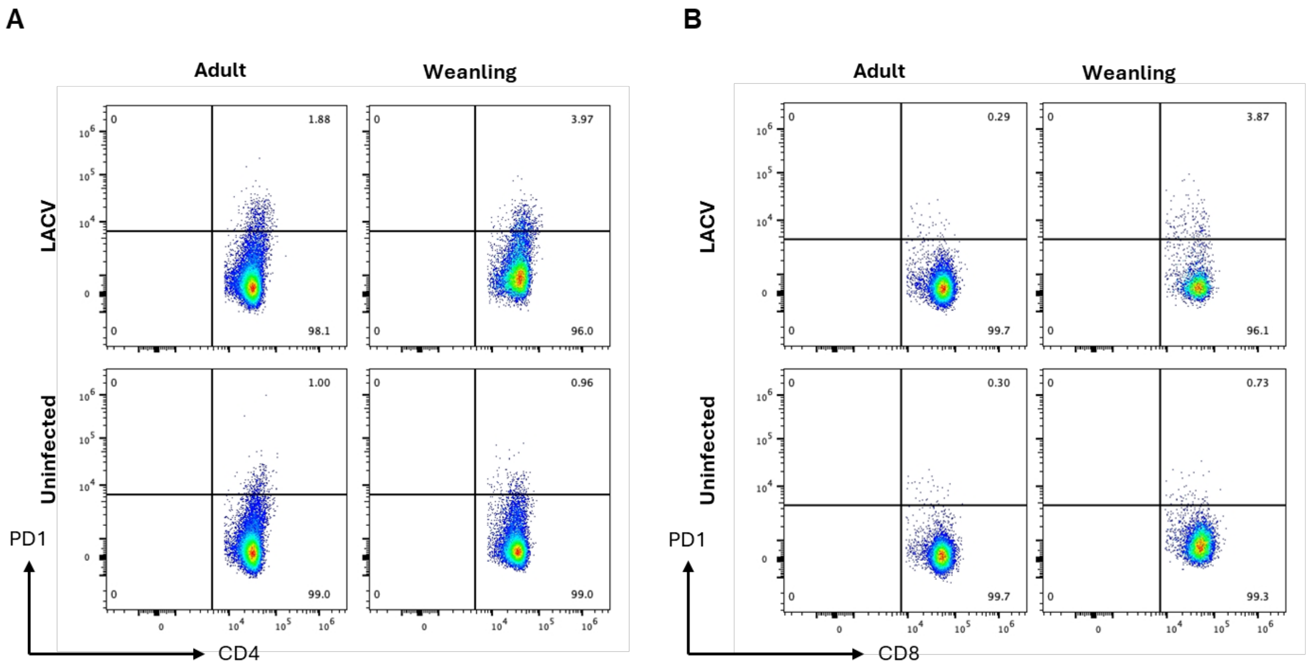


**Figure S1.** Representative flow cytometry plots showing gating of PD-1–positive $CD4^+$ (A) and $CD8^+$ T cells (B) from uninfected and LACV-infected mice across age groups. Quadrant gates indicate the frequency of $PD\text{-}1^+$ cells. Plots are representative of samples analyzed in each experimental group.

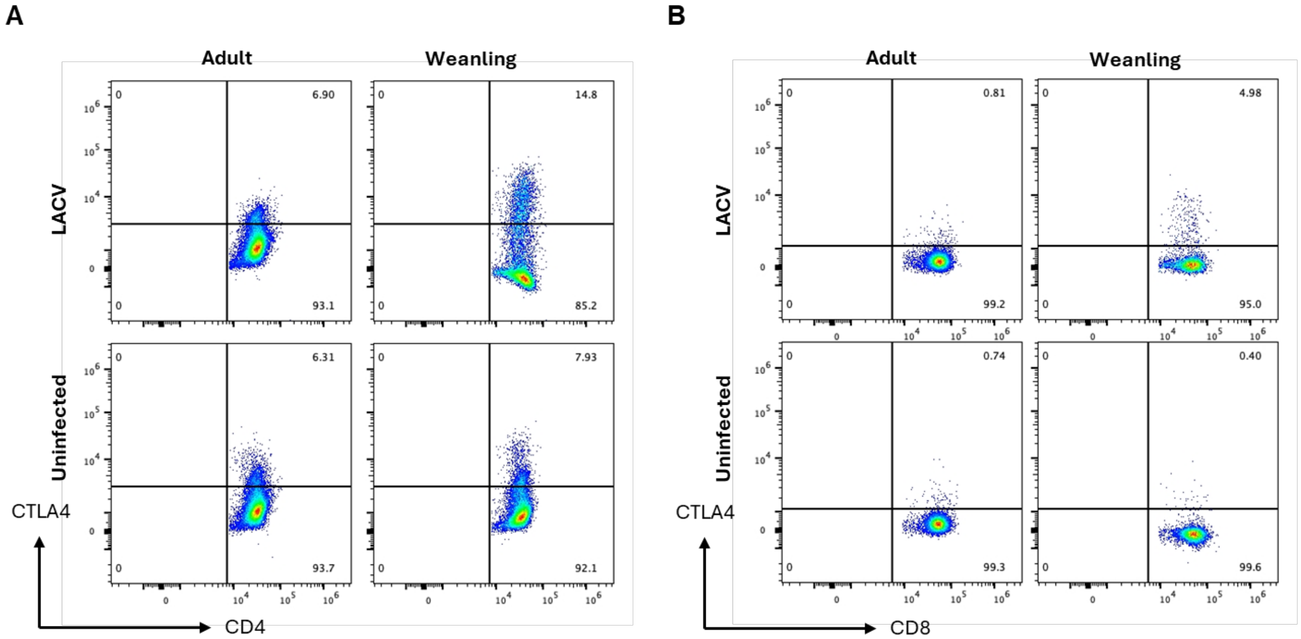


**Figure S2.** Representative flow cytometry plots showing gating of CTLA4-positive $CD4^+$ (A) and $CD8^+$ T cells (B) from uninfected and LACV-infected mice across age groups. Quadrant gates indicate the frequency of $CTLA4^+$ cells. Plots are representative of samples analyzed in each experimental group.

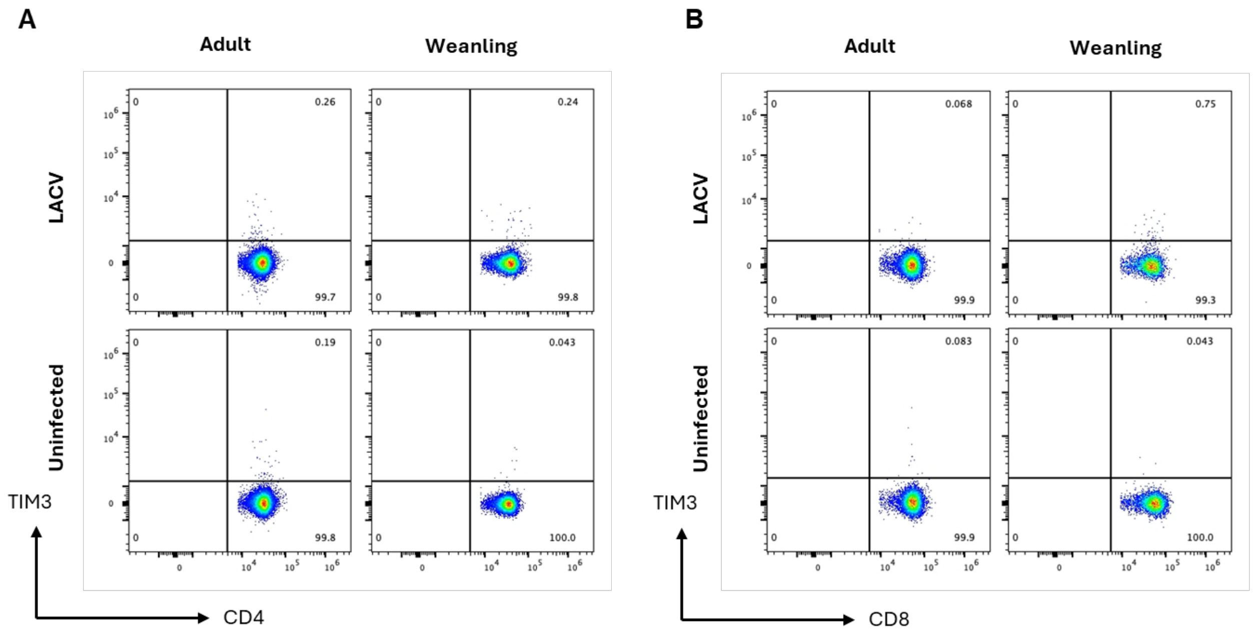


**Figure S3.** Representative flow cytometry plots showing gating of TIM3-positive $CD4^+$ (A) and $CD8^+$ T cells (B) from uninfected and LACV-infected mice across age groups. Quadrant gates indicate the frequency of $TIM3^+$ cells. Plots are representative of samples analyzed in each experimental group.

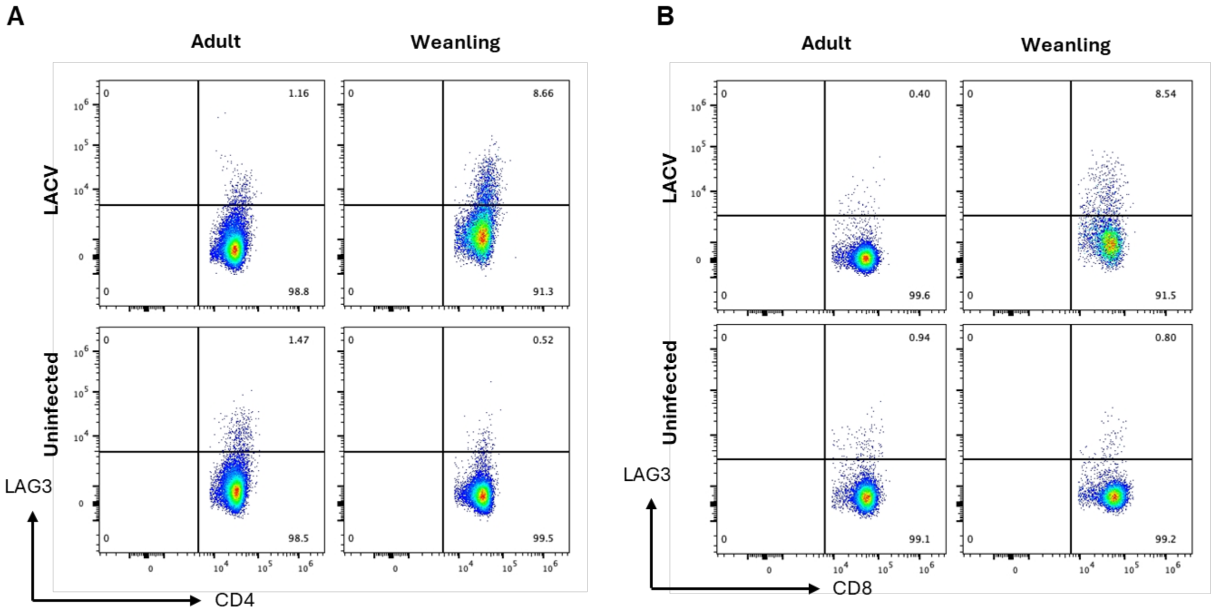


**Figure S4.** Representative flow cytometry plots showing gating of LAG3-positive CD4$^+$ (A) and CD8$^+$ T cells (B) from uninfected and LACV-infected mice across age groups. Quadrant gates indicate the frequency of LAG3$^+$ cells. Plots are representative of samples analyzed in each experimental group.

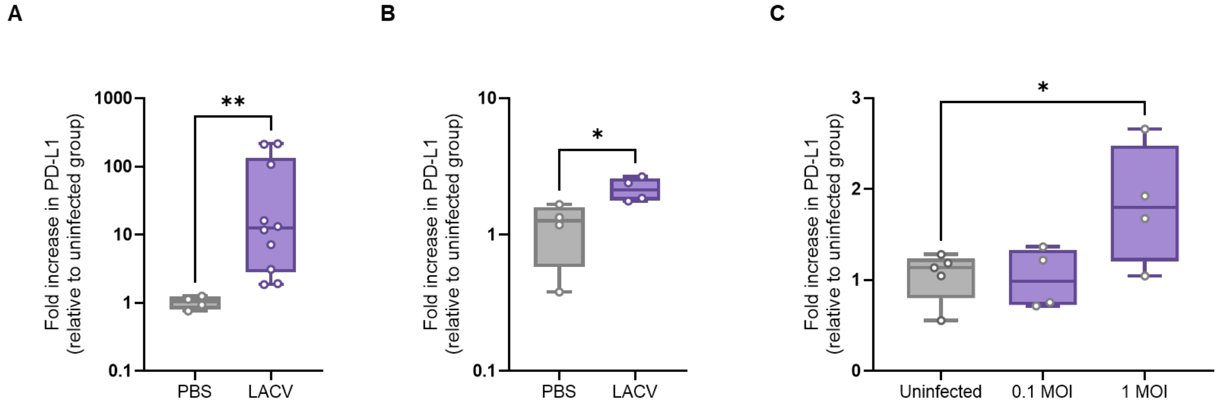


**Figure S5.** A) PD-L1 expression in the brain and (B) spleen of weanling mice following LACV infection compared to uninfected controls, as assessed by qPCR at 6 dpi. Each point represents an individual mouse, with box plots indicating fold change relative to uninfected mice within each group. (C) PD-L1 expression in mouse fibroblasts following LACV infection in vitro with 0.1 and 10 MOI compared to uninfected cells, as assessed by qPCR at 3 dpi. Statistical significance was determined using a Mann-Whitney test, $*p < 0.05$, $**p < 0.01$, $***p < 0.001$, $****p < 0.0001$. (n = 4-10 per group; data are representative of 2 independent experiments).

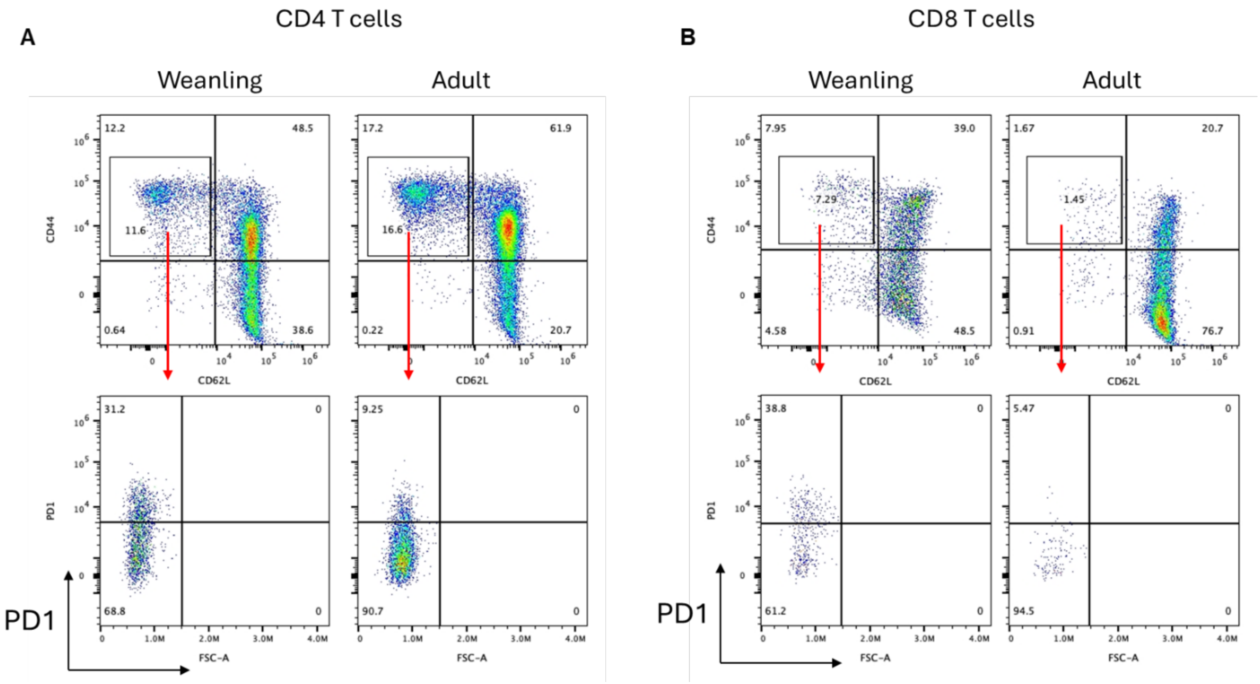


**Figure S6.** (A) Representative flow cytometry plots of CD44 versus CD62L expression on $CD4^+$ T cells and (B) $CD8^+$ T cells from weanling and adult mice following LACV infection at 6 dpi. Effector T cells were defined as $CD44^{hi}$ $CD62L^{lo}$ populations (top panels), and PD-1 expression was subsequently assessed within the effector subset (bottom panels). Arrows indicate the gating strategy used to identify effector T cells. Plots are representative of samples analyzed in each group.

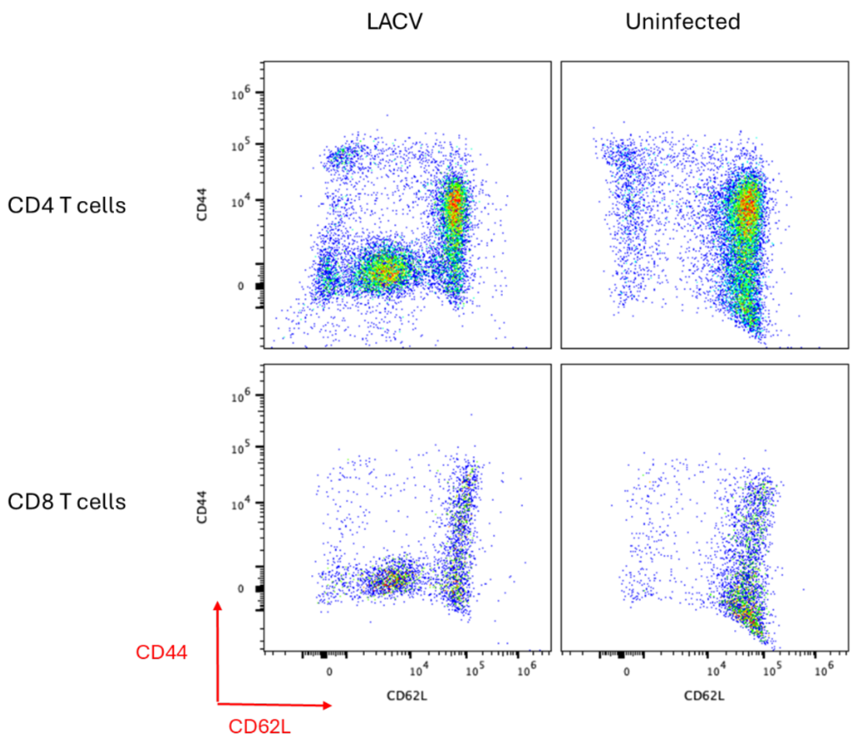


**Figure S7.** Representative flow cytometry plots of CD44 versus CD62L expression on $CD4^+$ (top panels) and $CD8^+$ (bottom panels) T cells from weanling mice that were either infected with LACV (left) or uninfected (right) at 6 dpi. Plots illustrate the distribution of naïve, central memory, and effector populations, with a notable accumulation of $CD44^-CD62L^-$ (double-negative) T cells following infection. Plots are representative of samples analyzed in each group.

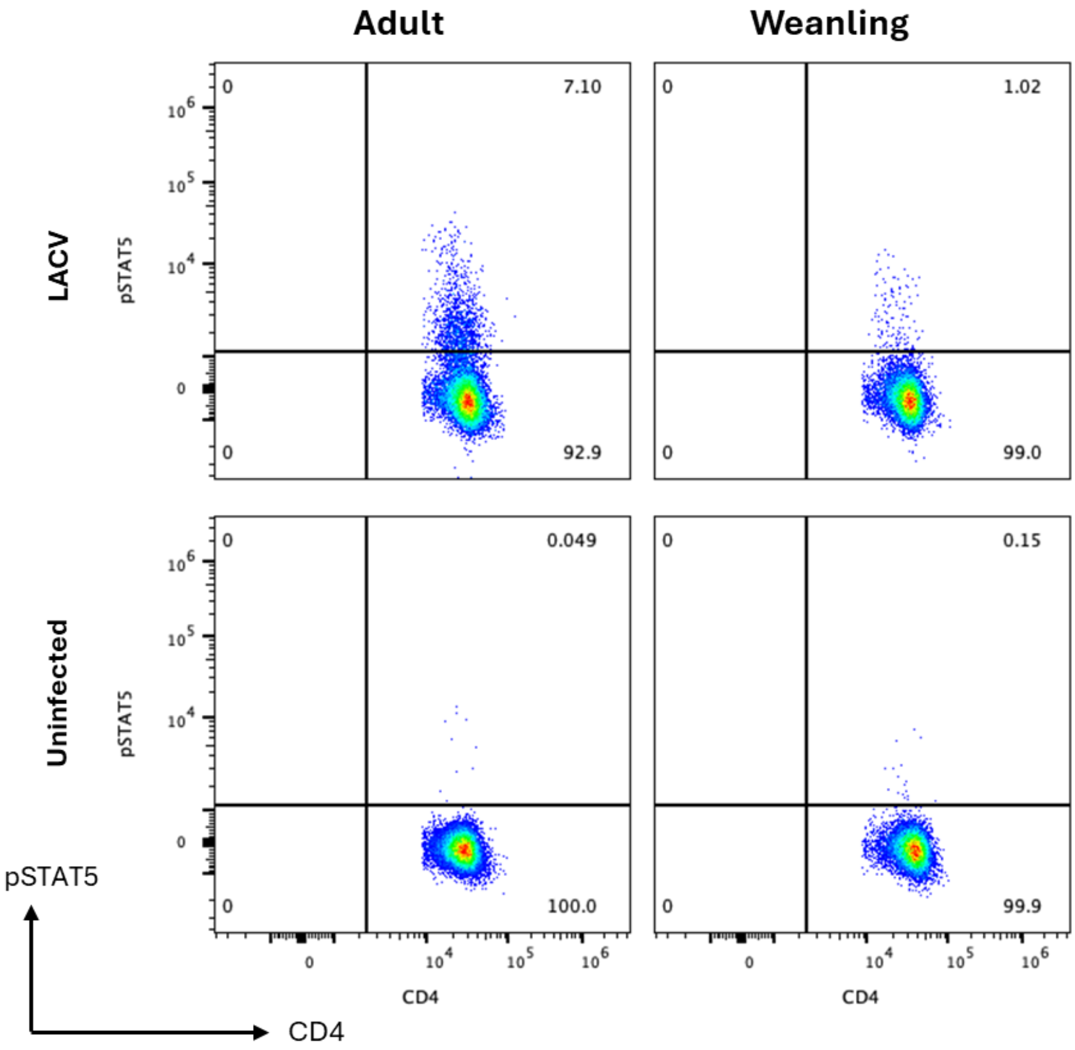


**Figure S8.** Representative flow cytometry plots of pSTAT5 expression in CD4⁺ T cells from weanling and adult mice. Top panels show LACV-infected mice and bottom panels show uninfected controls at 6 dpi. Quadrant gates indicate the frequency of STAT5⁺ cells within the CD4⁺ T cell population. Plots are representative of samples analyzed in each group.

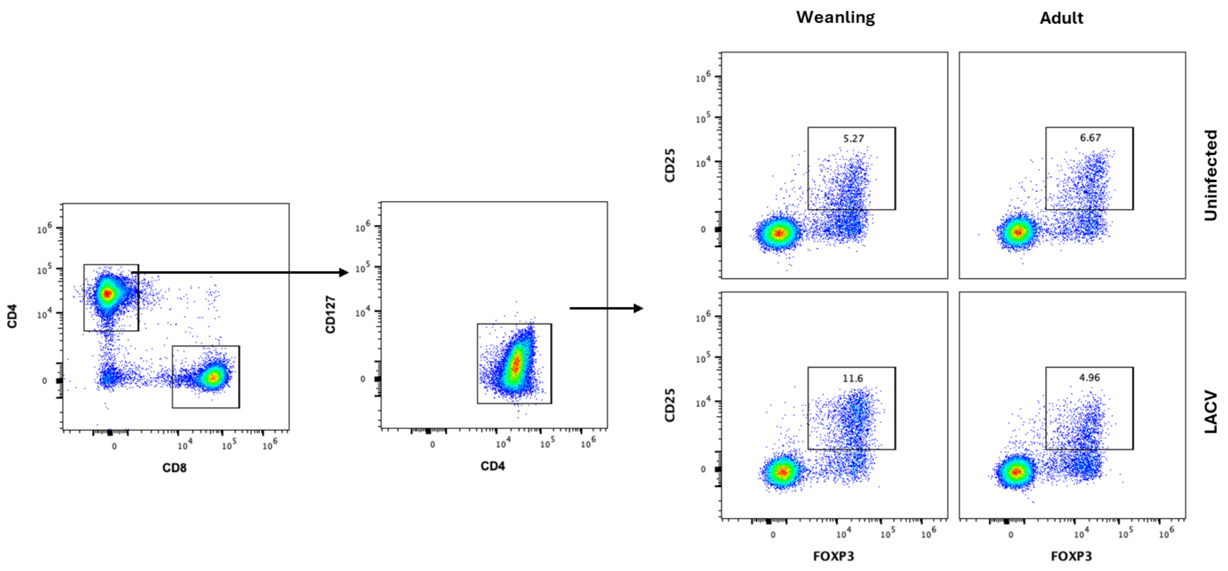


**Figure S9.** Representative flow cytometry plots showing the gating strategy used to identify CD4$^+$ regulatory T cells (Tregs). Cells were first gated on CD4$^+$ T cells, followed by selection of CD127$^{lo/-}$ cells, and subsequently CD25$^+$FoxP3$^+$ Tregs. Frequencies of Tregs are shown for weanling and adult mice under uninfected (top panels) and LACV-infected (bottom panels) conditions at 6 dpi. Plots are representative of samples analyzed in each group.

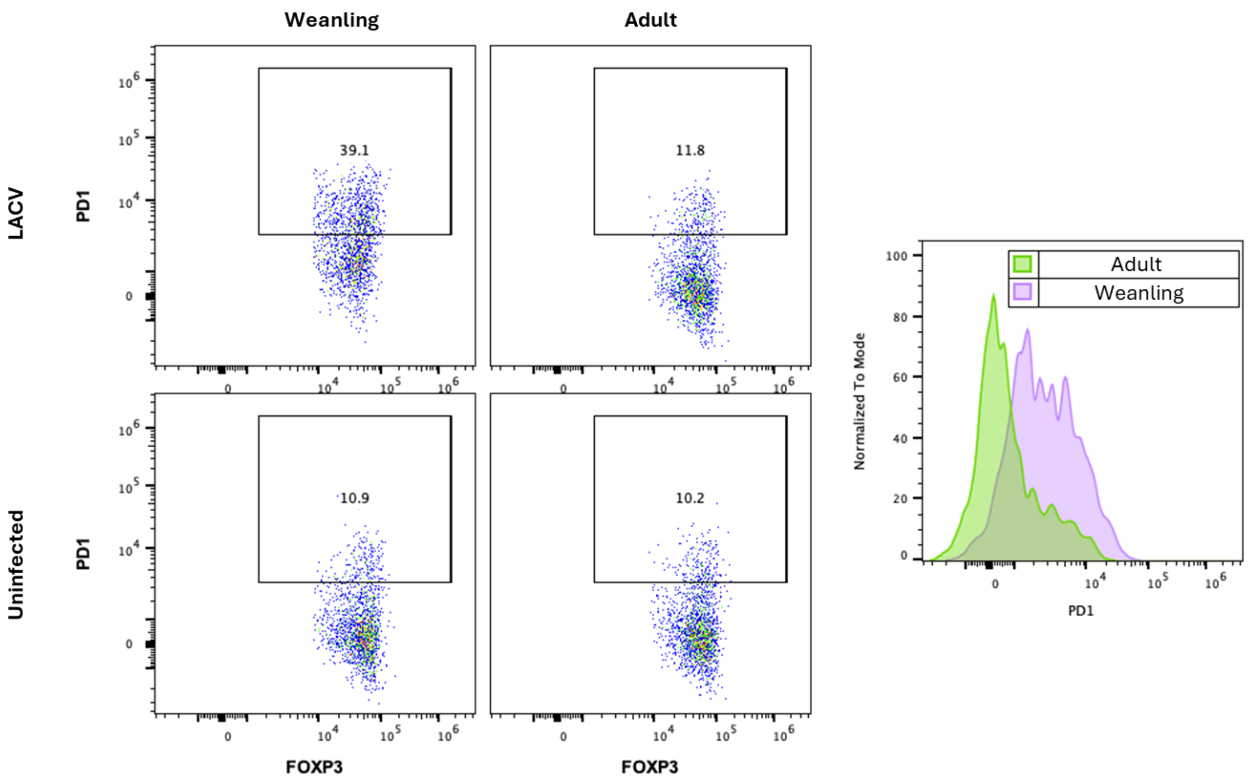


**Figure S10.** Representative flow cytometry plots showing PD-1 expression on CD4⁺ regulatory T cells (Tregs) from weanling and adult mice under LACV-infected (top panels) and uninfected (bottom panels) conditions at 6 dpi. Quadrant gates indicate the frequency of PD-1⁺ Tregs within each group. (Right) Histogram overlay of PD-1 expression on Tregs from infected weanling and adult mice. Plots are representative of samples analyzed in each group.

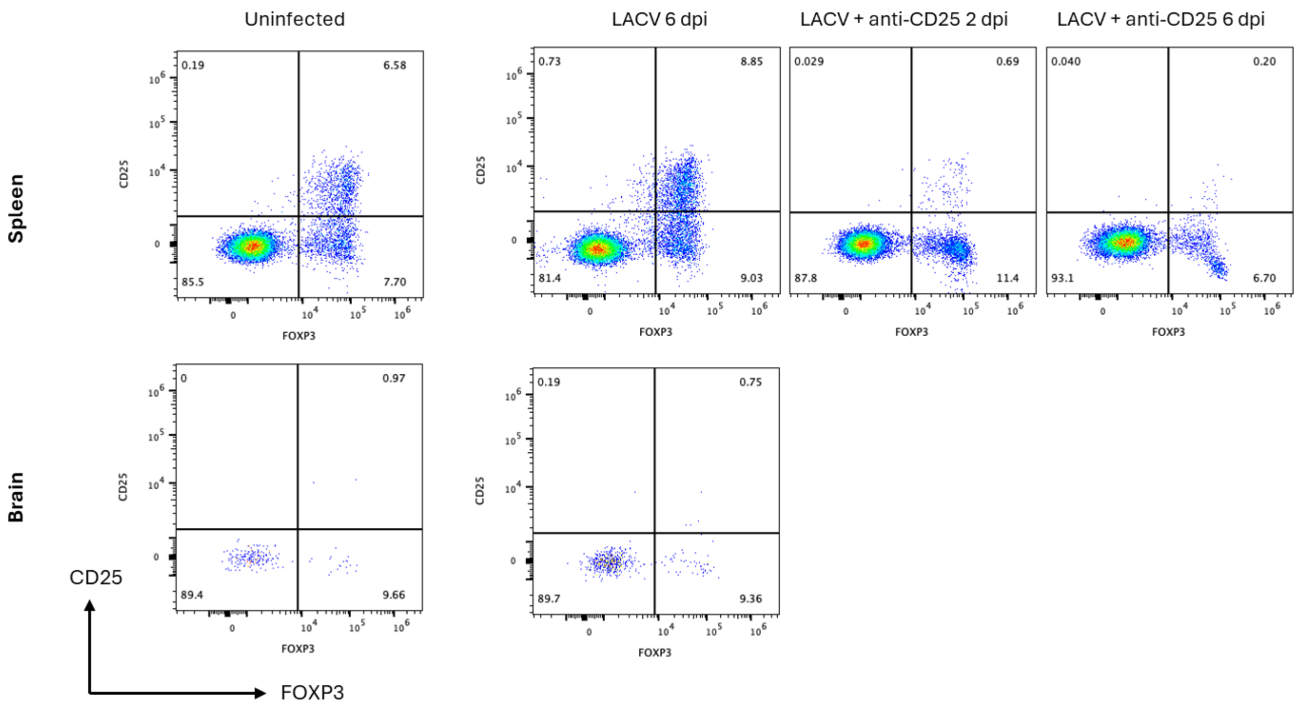


**Figure S11.** Representative flow cytometry plots showing frequencies of CD4$^+$ regulatory T cells (Tregs) in the spleen (top panels) and brain (bottom panels) from uninfected and LACV-infected mice. Treg depletion efficiency is shown in the spleen at 2 and 6 dpi following depletion treatment. Quadrant gates indicate the frequency of Tregs within each sample. Plots are representative of samples analyzed in each group.
